# Internal state dynamically gates task-specific attractor dynamics in prefrontal cortex

**DOI:** 10.64898/2026.05.20.726585

**Authors:** Yuma Osako, Timothy J. Buschman, Mriganka Sur

**Author notes:** Correspondence (Y.O.), (T.J.B), (M.S.).

## Abstract

Internal states such as motivation and task engagement influence cognitive functions. Working memory, which maintains information over time, is an essential component of cognition and is modulated by motivation. Here, we show motivational states modulated attractor dynamics that supported working memory. Combining population recordings from mouse medial prefrontal cortex (mPFC) with data-constrained recurrent neural network (RNN) modeling, we found task engagement selectively modulated attractor dynamics within a memory-maintenance subspace, while stimulus-evoked responses remained intact. Reverse-engineering the RNNs revealed that task engagement reorganized the dynamical landscape by stabilizing memory-specific attractors. Specifically, task engagement modulated interactions between neurons to change the attractor dynamics. Finally, gradual changes in behavioral engagement were predicted by continuous modulation of attractor geometry in RNNs and mPFC. Together, these results suggest that internal state modulate working memory function by controlling the dynamical regime of mPFC circuits, providing a mechanistic link between internal state, neural dynamics, and cognitive function.

## Introduction

Cognition is profoundly state-dependent. Our brain can perform vastly different computations, or perform the same computation with variable fidelity, depending on internal states such as attention^1^, reward expectation^2,3^, task engagement^4^, or arousal^5,6^. These internal states dynamically modulate neuronal activity, shaping how information is encoded, maintained, and transformed across cognitive processes^7^. One of the central challenges in neuroscience is to understand how internal states dynamically reconfigure neuronal dynamics to engage in specific cognitive functions. However, it remains unclear how latent variables, such as task engagement or motivational fluctuations influence the neural substrates of complex cognitive functions.

Working memory, the capacity to temporarily maintain and manipulate information in the absence of sensory input, is one cognitive function that is particularly sensitive to cognitive control^8^. It is an active process, thought to rely, at least in part, on persistent neuronal activity in the prefrontal cortex (PFC) to form a stable representation of task-relevant information^9–19^. Theoretical work suggests that such persistent activity can be explained through attractor dynamics, in which network activity evolves toward and stabilizes around specific patterns of neuronal activity (known as attractors)^20–25^. However, attractor dynamics are not immutable, rather they are often modulated across different contexts and can exhibit trial-by-trial variability even within the same task^18,26^. Many previous studies have shown that single neuron and population-level activity can vary depending on context^27–29^ and show subtle fluctuations associated with internal states^6,30–32^, such as task engagement^27^. These observations suggest that internal states dynamically regulate neural circuits and, in turn, reshape attractor dynamics in the brain^21,33^. However, how internal states influence the emergence, stability, and reconfiguration of attractors remains largely unknown.

Here, we address this question by combining population recordings from the mouse medial PFC (mPFC)^34^ with recurrent neural network (RNN) modeling. By comparing neuronal dynamics during active task engagement and passive stimulus exposure, we show that task engagement selectively modulates memory-related subspaces. Data-constrained RNNs reveal bistable attractors that support memory maintenance during the active task engagement, whereas these collapse into a single attractor during passive conditions. This bifurcation can be explained by context-dependent modulation of neuronal interactions, as supported by both the model and mPFC data. Furthermore, we found that gradual fluctuations in task motivation during a recording session in mPFC correlated with graded shifts in attractor separation in the model, directly linking internal states, network dynamics, and behavioral performance in the brain. Together, these findings establish a mechanistic framework for how internal states dynamically gate working memory by reshaping attractor dynamics in cortical circuits.

## Results

### The delayed match-to-sample with delayed report task

Mice were trained on a DMS-dr task, in which they were asked to report whether two auditory stimuli were the same or different^34^. The stimuli were pure-tone 0.3s auditory stimuli with either a high (H, 14kHz) or low (L, 3kHz) frequency. The first and second stimuli were separated by a 1s memory delay. After the second stimulus, there was another 1s delay before the animal reported whether the two stimuli matched, by licking a reward spout, or did not match, by refraining from licking. Correct licks (hits) were rewarded with water, while incorrect licks to non-match stimuli (false alarm) resulted in an air-puff and a 7s-timeout (**Figure 1A**). Trials in which the mice refrained from licking after matched stimuli (miss) or after non-match stimuli (correct reject) were not reinforced. To determine how task engagement influenced network dynamics, mice also performed a passive listening block after performing the DMS-dr task during which the reward spout was not presented and so the animals could not respond (**Figure 1B**). As expected, mice did not exhibit licking behavior during the passive condition (**Figure S1A–S1C**), whereas robust licking was observed during the task condition after the spout was reached around mouth.

**Figure 1.**
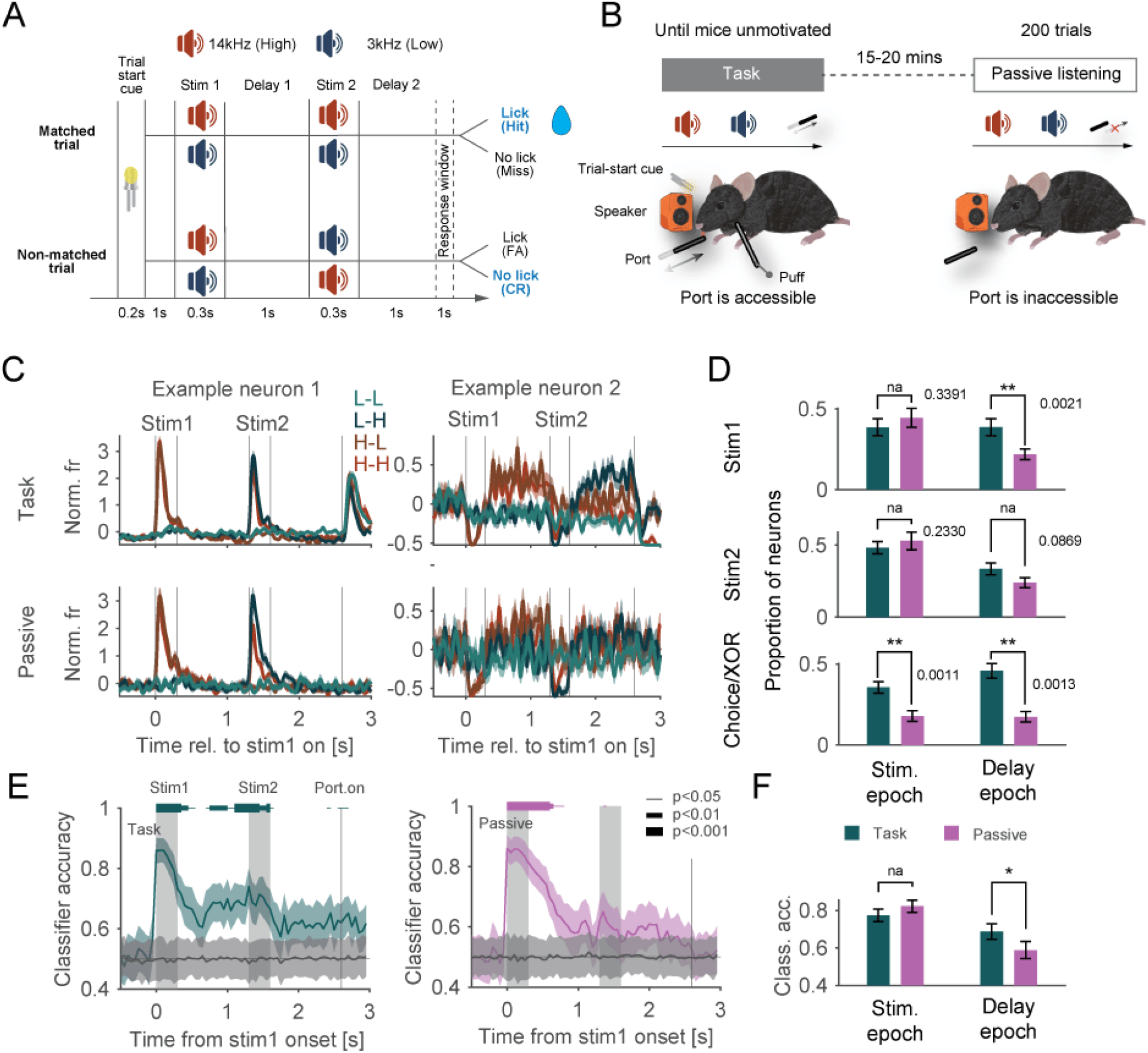
Neuronal activity during a DMS-dr task. **(A)** Timeline and trial types of the DMS-dr task. Licking responses were measured within a 1-s response window. **(B)** Experimental setup for active task engagement (left) and passive stimulus exposure (right). **(C)** Example neuronal activity during active and passive tasks, with colors indicating the four trial types. **(D)** Proportion of neurons encoding stimulus 1, stimulus 2, and choice/XOR during stimulus and delay epochs in active and passive tasks. **(E)** Time course of classifier accuracy for stimulus 1 in active (left) and passive (right) tasks. Horizontal bars above each plot indicate periods of above-chance classification (P < 0.05, 0.01, and 0.001 for thin, medium, and thick lines, respectively; bootstrap test). **(F)** Classifier accuracy for stimulus 1 during stimulus and delay epochs in active and passive tasks.

### Task engagement modulates delay-period activity in mPFC

We recorded neuronal activity from the medial prefrontal cortex (mPFC) of mice performing both the active DMS-dr task and during passive stimulus exposure (**Figure 1B**). Previously, we showed mPFC is causally engaged in this task^34^. We observed robust and selective responses to stimulus 1 during its presentation in both task and passive conditions (**Figure 1C**, Example neuron 1). However, the sustained representation of stimulus 1 during the delay period was attenuated in the passive condition compared to the DMR-dr task (**Figure 1C**, Example neuron 2). To quantify this, we assessed the selectivity of individual neurons for stimulus 1, stimulus 2, and matched/non-matched trials across different task epochs (see Methods, **Figure S2**). We found the proportion of neurons selective for stimulus 1 and stimulus 2 during stimulus presentation did not significantly differ between the task and passive conditions (**Figure 1D**; stimulus 1 selectivity during stimulus epoch: task = 38.22%, passive = 46.17%, p = 0.1699, paired t-test; stimulus 2 selectivity during stimulus epoch: task = 45.13%, passive = 52.74%, p = 0.1842). Accordingly, because no memory of the stimuli is required under passive conditions, activity during delay 2 period is expected to be comparable between task and passive conditions. In contrast, selectivity for the memory of stimulus 1 during the delay 1 was significantly reduced in the passive condition (**Figure 1D**; task = 37.36%, passive = 22.13%, p = 0.0011, paired t-test). Similarly, selectivity for the memory of the upcoming response was also significantly diminished in the passive condition during both the stimulus and delay epochs (**Figure 1D**; stimulus epoch: task = 35.98%, passive = 18.22%, p = 0.0017; delay epoch: task = 49.17%, passive = 16.69%, p = 6.24e-5). As the task did not require the animals to remember the second stimulus, selectivity for the second stimulus was reduced during the task and then even further reduced during the passive condition (stimulus 2 selectivity during delay epoch: task = 34.84%, passive = 24.36%, p = 0.1210). Together, these findings suggest that sensory encoding of stimuli in mPFC is preserved irrespective of task engagement but that the active maintenance of task-relevant information for working memory depended on task engagement.

To further assess how these selectivity differences manifest at the population level, we performed classification analysis. We trained logistic regression classifiers using activity from all neurons pooled across experiments (pseudo-population^35^) to decode the identity of stimulus 1 information (see Methods). Classifiers were fitted independently within 50-ms sliding time windows across the entire trial (**Figure 1E**). Consistent with single neuron analyses, stimulus 1 information could be decoded around stimulus presentation in both task and passive conditions (**Figure 1E**). Quantitatively, the average classification accuracy during the stimulus epoch did not differ significantly between conditions (**Figure 1F**, p>0.05, bootstrap test), whereas accuracy during the delay 1 epoch was significantly decreased in the passive condition compared to the task (task = 68.74%, passive = 58.84%, p<0.05, bootstrap test). Similar to the single-neuron analysis, classification accuracy for match/non-match was also significantly higher in the task compared to the passive condition (**Figure S3A-B**).

During the delay 2 period, mice are required to maintain match/non-match information to prepare a motor response. The attenuation of information in the passive condition therefore suggests that they either did not engage working memory or failed to integrate stimulus 2 with stimulus 1. These results indicate that task engagement enables the active maintenance of memories during delay periods in the mPFC, whereas the encoding of sensory stimuli remains largely stable regardless of task conditions.

### Task engagement selectively affects memory representations in recurrent neural networks

Our single-neuron and population-level analyses revealed that memory maintenance was modulated by task engagement. To gain insight into the underlying mechanisms, we trained recurrent neural networks (RNNs) to perform the DMS-dr task while reproducing the neuronal dynamics observed in the mPFC (**Figure 2A**). The RNNs received three types of noisy inputs: stimulus, fixation, and context inputs (**Figure 2A**). Stimulus inputs mimicked high and low tones in DMS-dr task. Fixation inputs signaled to the network when to respond. The context input indicated whether the network should respond based on the stimulus inputs (active task condition) or not (passive task condition; **Figures 2A** and **S4A**). To capture biological responses, all units in the RNN were trained to produce non-negative activity through the sigmoid activation function.

**Figure 2.**
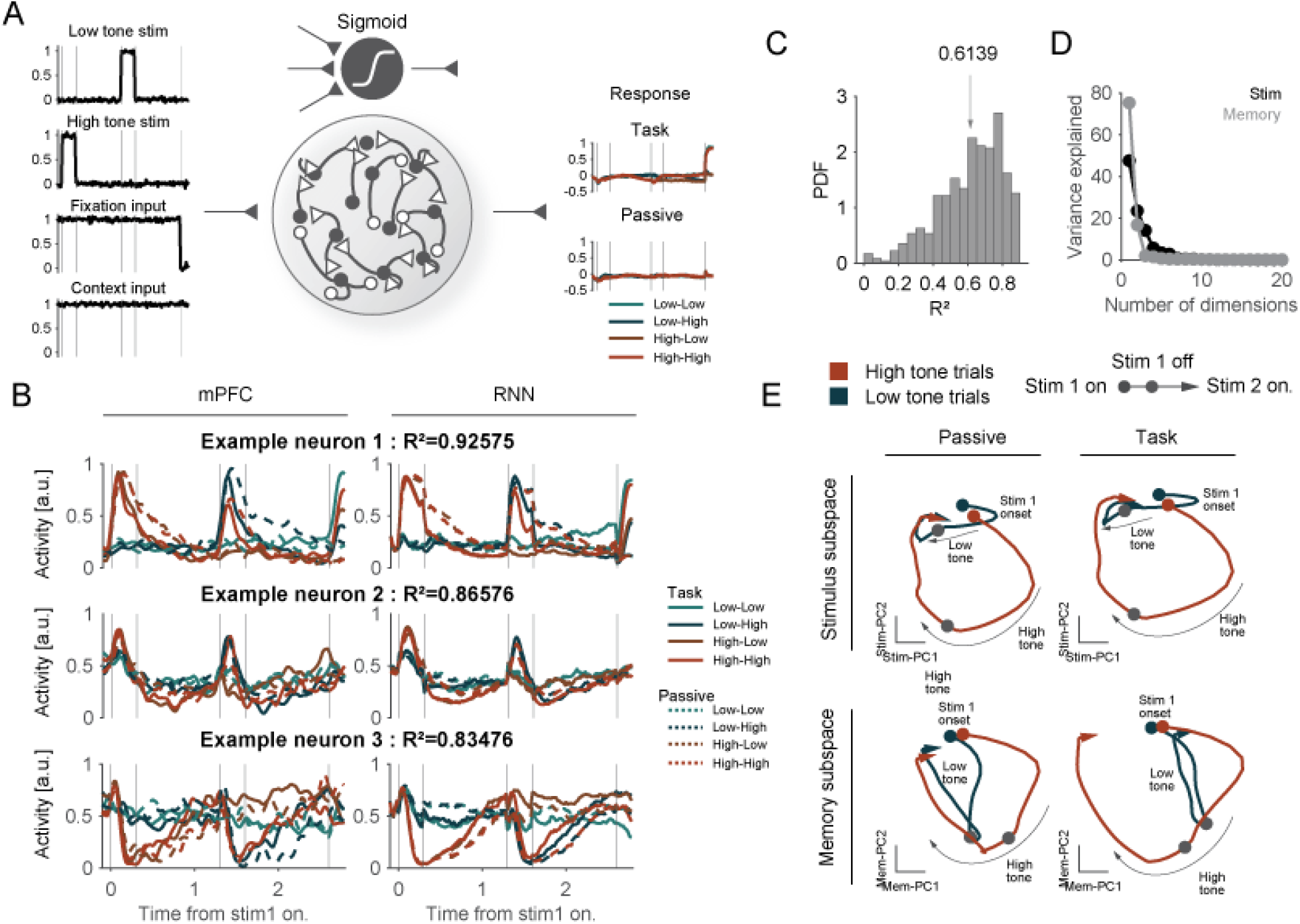
Data-constrained RNN robustly reproduce mPFC neuronal activity. **(A)** Schematic of a fully connected RNN architecture. The network receives stimulus, fixation, and context inputs, and projects to a behavioral output unit. All RNN units exhibit non-negative responses through a ReLU activation function. An RNN containing 443 units was trained to mimic neuronal activity during active and passive tasks. **(B)** Example RNN unit activities (right) trained to reproduce the PSTHs of neurons (left) across eight unique trial types. **(C)** Histogram of R^2^ values for all RNN units, computed by comparing PSTHs across eight trial types between RNN units and mPFC neurons. **(D)** Explained variance for the top 20 dimensions of the stimulus and memory subspaces. **(E)** Population activity projected onto stimulus and memory subspaces during active and passive tasks. Red and green lines indicate high- and low-tone trials, respectively.

The RNNs reliably reproduced the peri-stimulus time histograms (PSTHs) of mPFC neurons recorded during both task and passive conditions (**Figure 2B**), with individual units capturing a significant fraction of neural variance across time and task condition (**Figure 2C**, *Mean R*^2^ = 0.6139). The variance explained was comparable between task and passive conditions (**Figure S4B**, 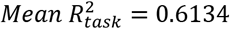 and 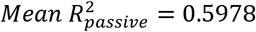, *p* = 1.0 , Two-sample Kolmogorov-Smirnov test), indicating that the RNN reliably reproduced the mPFC activity for both the active and passive conditions.

To examine how population-level representations evolved across conditions, we used the first two principal components (PCs) of the RNN activity to define a stimulus subspace during the stimulus presentation (0–0.5 s from stimulus 1 onset) and to define a memory subspace during the delay (0.8–1.3 s; **Figure 2D**). Both subspaces captured over 70% of the total variance of neural activity in their epoch. This provides a reference for the fraction of variance captured in visualizations for subsequent analysis. We next visualized how the population trajectory onto these top two subspaces evolved during stimulus 1 and delay 1 epoch (**Figure 2E**, -0.1-1.3s from stimulus 1 onset). Projecting the population activity onto the stimulus subspace revealed distinct trajectories for high and low tone trials that diverged during the stimulus presentation and then reconverged during the delay period (**Figure 2E**, top row). These dynamics were similar for both the active and passive task conditions. In the memory subspace, neural activity remained separated throughout the delay in the active task condition (reflecting the sustained memory) but gradually converged in the passive task condition (**Figure 2E**, bottom row). This selective modulation of the memory subspace by context input was consistent with the mPFC data (**Figure 1C-1F**).

To understand how the task context influence memory dynamics, we systematically interpolated the amplitude of the context input between the passive (0) and active (1) task conditions in increments of 0.05, and traced trajectories across conditions (**Figure 3A**). While trajectories in the stimulus subspace remained largely invariant across interpolated context input, those in the memory subspace exhibited gradual divergence during the delay epoch as a function of amplitude of the context input. To quantify this divergence, we computed the Euclidean distance between high and low tone trajectories at each context level (**Figure 3B**). This analysis revealed that Euclidean distance in the stimulus subspace was largely unaffected by context (**Figure 3B**, left) but that increasing the context signal increased the separation between high and low tones in the memory subspace (**Figure 3B**, right).

**Figure 3.**
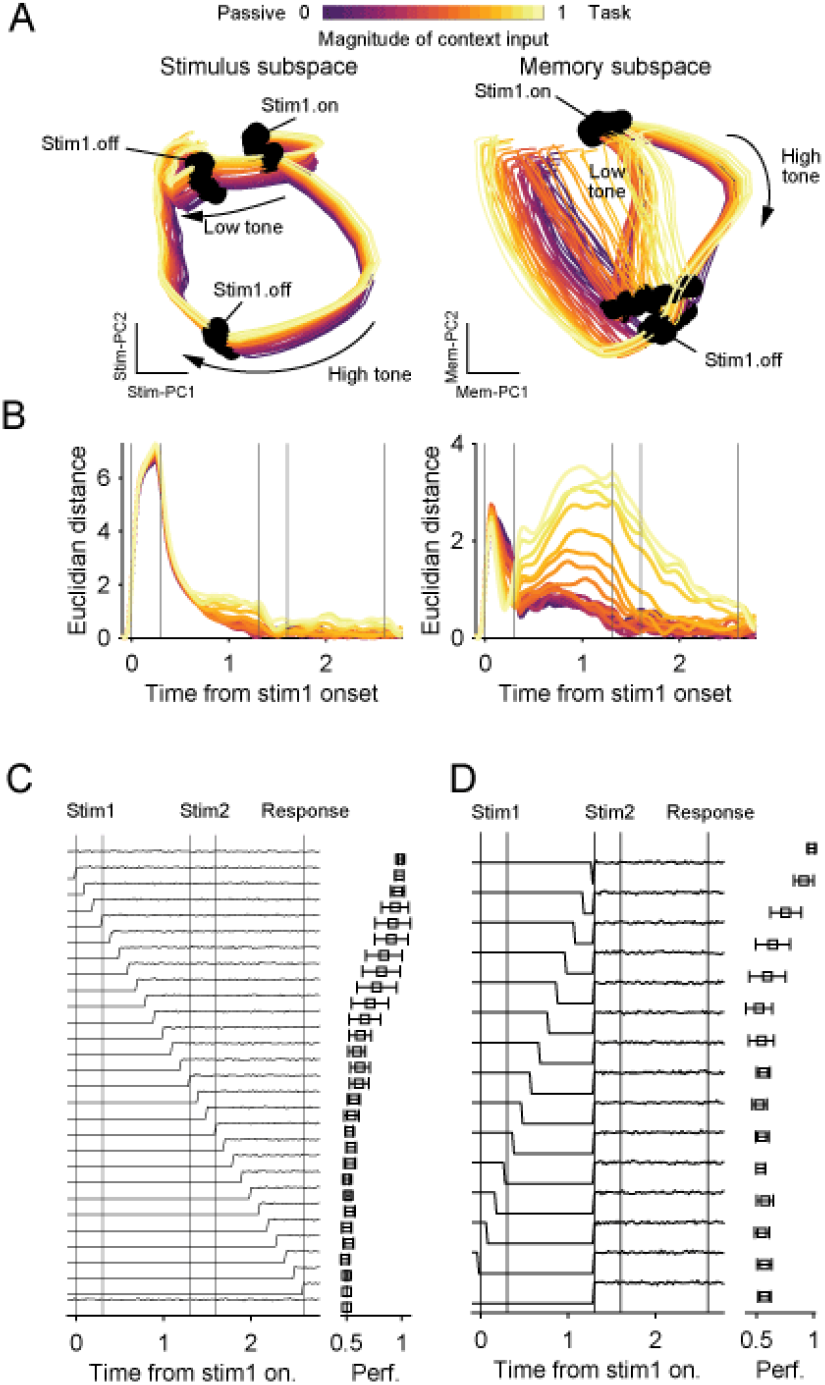
Context input specifically modulates memory maintenance dynamics. **(A)** (top) Interpolation between inputs for passive (input = 0) and active (input = 1) tasks. (Bottom) Population activity projected onto the stimulus and memory subspaces for each magnitude of context input. **(B)** Euclidean distance between low- and high-tone trials in the stimulus and memory subspaces. **(C)** RNN performance under different types of context inputs. Context inputs were shifted from 0 to 1 at different timings. **(D)** Same as (C), but with different shapes of context inputs.

To understand when context influenced circuit dynamics, we systematically varied when context was provided during the trial of the active task (**Figure 3C**). Network behavior was not disrupted by the absence of contextual input before the delay epoch. However, performance gradually degraded when the contextual input was set to 0 after the stimulus epoch. Crucially, even if the context input was switched to 1 during the latter half of delay epoch (around 0.8s for stimulus 1 onset), network performance remained significantly impaired. This shows memory maintenance requires context input from the middle of delay period but that context input is not required for encoding the stimulus. We also confirmed that context input during delay 2 period is required for accurate behavior (**Figure S4C**).

Next, we tested whether memory maintenance required continuous context input throughout the delay period or if a brief input at the onset of the delay period was sufficient to drive the network into memory subspace. To discriminate these hypotheses, we switched the context input from 1 to 0 before the onset of the stimulus 2, and then switched back to 1 upon its presentation (**Figure 3D**). We found that network performance significantly degraded when the context input was 0 at any point during the delay period except for last 100ms (**Figure 3D**, top two), indicating that continuous input is required to sustain memory-related dynamics. The RNNs could still perform above chance level when the context input was 0 during the late delay period, likely due to the time constant (200ms). Together, these results suggest that sustained context input during the delay period is necessary for correct behavioral performance and acts selectively on the memory-maintenance subspace.

### Context input drives bistable attractor dynamics

We next reverse engineered the RNNs to uncover the mechanism by which context input enables memory maintenance. During delay 1 in the active task condition, dynamics in the memory subspace pulled neural representations into one of two stable attractors corresponding to high and low tone trials, separated by a saddle point (**Figure 4A**, left; **Figure S5A**). In contrast, in the passive condition, population dynamics converged onto a single attractor (**Figure 4A**, right; **Figure S5B**), showing that the dynamics of the network cannot sustain two stimulus representations. By systematically interpolating the context input from passive to active task conditions, we observed an emergence of the two attractors and a saddle point during the intermediate stages of interpolation (around 0.7 of context input, **Figure 4B**). As the contextual input incrementally increased, it drove the two attractors further apart, concomitantly increasing the distance between the low- and high-tone trajectories (**Figure 4C**, top). This bifurcation pattern was consistently observed across additional networks (**Figure S5C**, 8 of 10 networks). Moreover, this separation of attractors was positively correlated with the RNNs behavioral performance (**Figure 4C**, bottom). These results suggest that the context input tunes the dynamics to create distinct attractors (corresponding to low and high tone) as the system transitions from the passive to active task condition.

**Figure 4.**
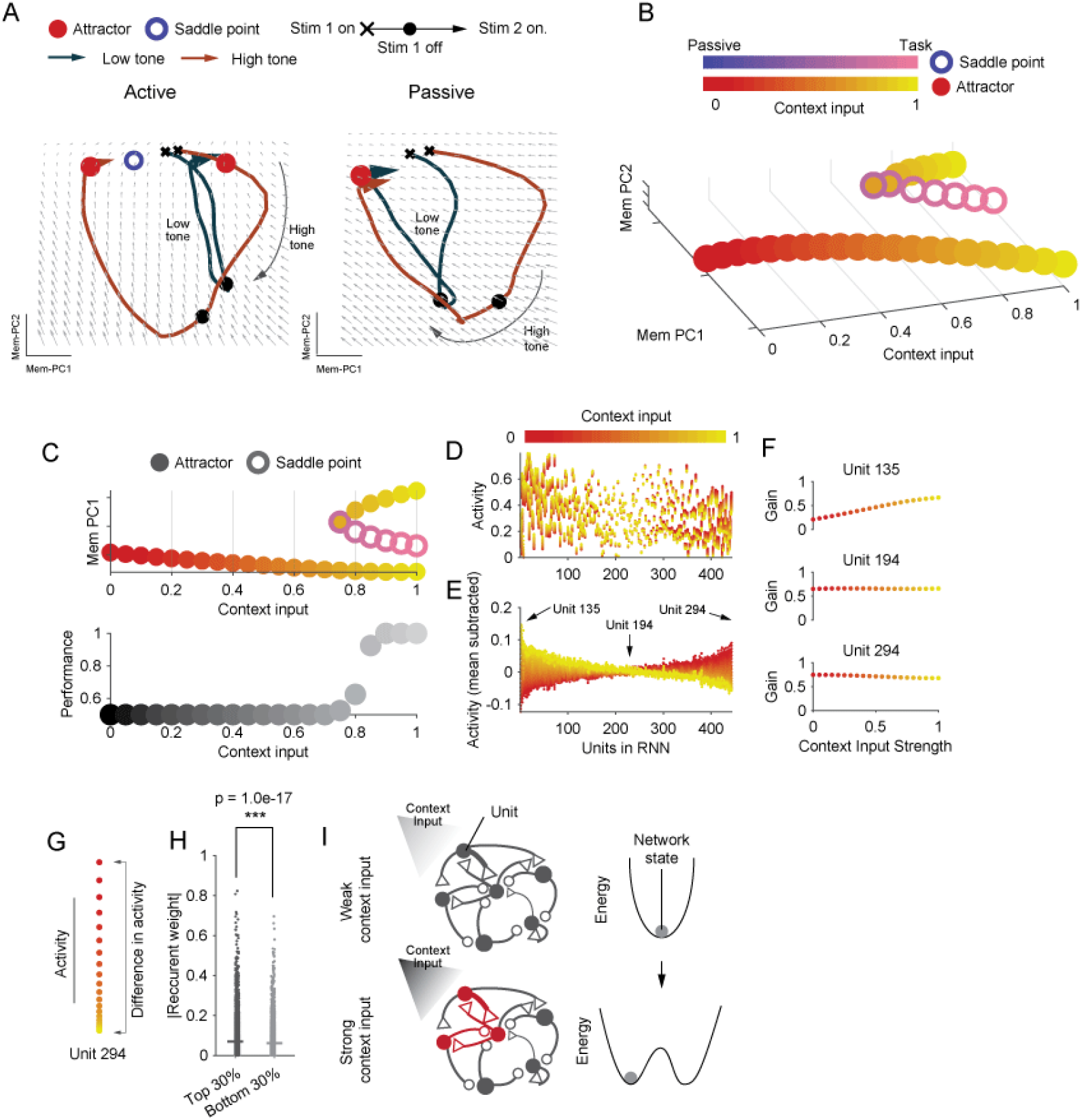
Gradual reorganization of attractors with context inputs. **(A)** Attractors (red points) and saddle points (blue circles) in the memory subspace during active (left) and passive (right) tasks. Lines represent population activity trajectories from stimulus 1 onset to stimulus 2 onset. **(B)** Attractors and saddle points during interpolation between contextual input strengths corresponding to the passive (input = 0) and active task (input = 1) conditions, visualized in the memory subspace. Dots and circles denote attractors and saddle points, respectively, with color indicating the magnitude of contextual input. **(C)**(Top) Same as in B, but projected onto the first principal component (PC) of the memory subspace. (Bottom) RNN performance during interpolation between passive and active inputs. **(D)** RNN units were sorted according to their activity differences between the passive and active conditions. Unit activity in the vicinity of the attractor is plotted as a function of contextual input strength. **(E)**Same as (D), after mean subtraction across units. **(F)** Gain modulation of example RNN units. Increasing context input drives units toward high gain (unit 135) or low gain (unit 294), whereas other units remain largely insensitive (unit 194). **(G)** Schematic illustrating context input dependent variability in unit slope. **(H)** Absolute recurrent connectivity weights between the top and bottom 30% of units exhibiting the largest differences in activity between the passive and active conditions. **(I)** Mechanism of bifurcation revealed by the data-constrained RNN. In the passive condition, network dynamics exhibited a single baseline attractor. Introduction of context input induced a bifurcation, giving rise to two attractors aligned with low- and high-tone trajectories, separated by an intermediate saddle point. Increasing context input strength progressively reshaped the dynamical landscape, switching the network into a distinct dynamical phase supporting stimulus-specific memory states.

We next examined how context input modulates neuronal activity in the RNN. Context input induced highly heterogeneous shifts in unit activity in the vicinity of the attractor across the population (**Figure 4D** for raw activity; **Figure 4E** for mean subtracted). Furthermore, gain modulation was unit-specific (**Figures 4F** and **S5D**), indicating that context input does not impose a uniform bias but rather differentially modulates neuronal activity in the network. One hypothesis for why there is a heterogenous control input is that it specifically targets circuits that sustain memories. To test this, we asked whether units strongly modulated by context input were preferentially interconnected. Units were sorted according to the magnitude of activity change between the passive and task conditions (**Figure 4G**) and divided into the top and bottom 30% of the distribution. Recurrent connectivity among the top 30% of units was significantly stronger than that among the bottom 30%, indicating that context-sensitive units form a more strongly interconnected subnetwork (**Figure 4H**). Similar results were obtained when sorting units by differences in gain across passive and task conditions (top and bottom 10–40%; **Figure S5E**). Taken together, these results suggest that context input selectively engages a subset of highly context-sensitive units and their recurrent interactions, driving a transition in the network’s dynamical phase to give rise to the observed bifurcation (**Figure 4I**).

### Context-dependent modulation of memory representations by behavioral state

Our analyses thus far indicate that gradual changes in context input dynamically shape attractor dynamics in the RNN. Although we initially modeled contextual input as a binary variable distinguishing active and passive task conditions, we hypothesized that neural activity may also reflect more subtle, behaviorally relevant fluctuations, such as changes in the animal’s motivational state. To test this possibility, we divided trials into octiles based on task progression and computed hit rates for each bin. Hit rates showed a strong negative correlation with trial progression (r = −0.844, p < 0.001; **Figure 5A**), suggesting that the animals’ motivation decreased as the session went on. We then asked whether this behavioral change was accompanied by corresponding alterations in neural dynamics. To this end, we quantified the Euclidean distance between population trajectories evoked by high- and low-tone stimuli within the stimulus and memory subspaces. While stimulus-epoch activity remained stable across trial progression, delay-period activity within the memory subspace fluctuated as a function of trial progression (**Figure 5B**, top). Importantly, these trial-by-trial fluctuations were strongly correlated between mPFC recordings and the RNN selectively during the delay epoch in the memory subspace (**Figure 5C**). No significant correlations were observed during the stimulus epoch in either subspace (stimulus epoch: r = 0.1328 and 0.4030, p = 0.6873 and 0.1870 for stimulus and memory subspaces, respectively), whereas there was a significant correlation during the delay epoch in the memory subspace (delay epoch: r = −0.4796 and 0.6638, p = 0.1169 and 0.025 for stimulus and memory subspaces, respectively; permutation test). This effect was consistent across all trained networks (**Figure 5D**). Together, these results suggest that gradual changes in behavioral state during the session likely reflected fluctuations in motivation, which, in turn, modulated population dynamics during memory maintenance. This pattern closely parallels how graded context input modulates attractor dynamics in the RNN, supporting the idea that memory representations in mPFC are not static but are continuously reshaped by latent behavioral variables that modulate the underlying dynamical landscape.

**Figure 5.**
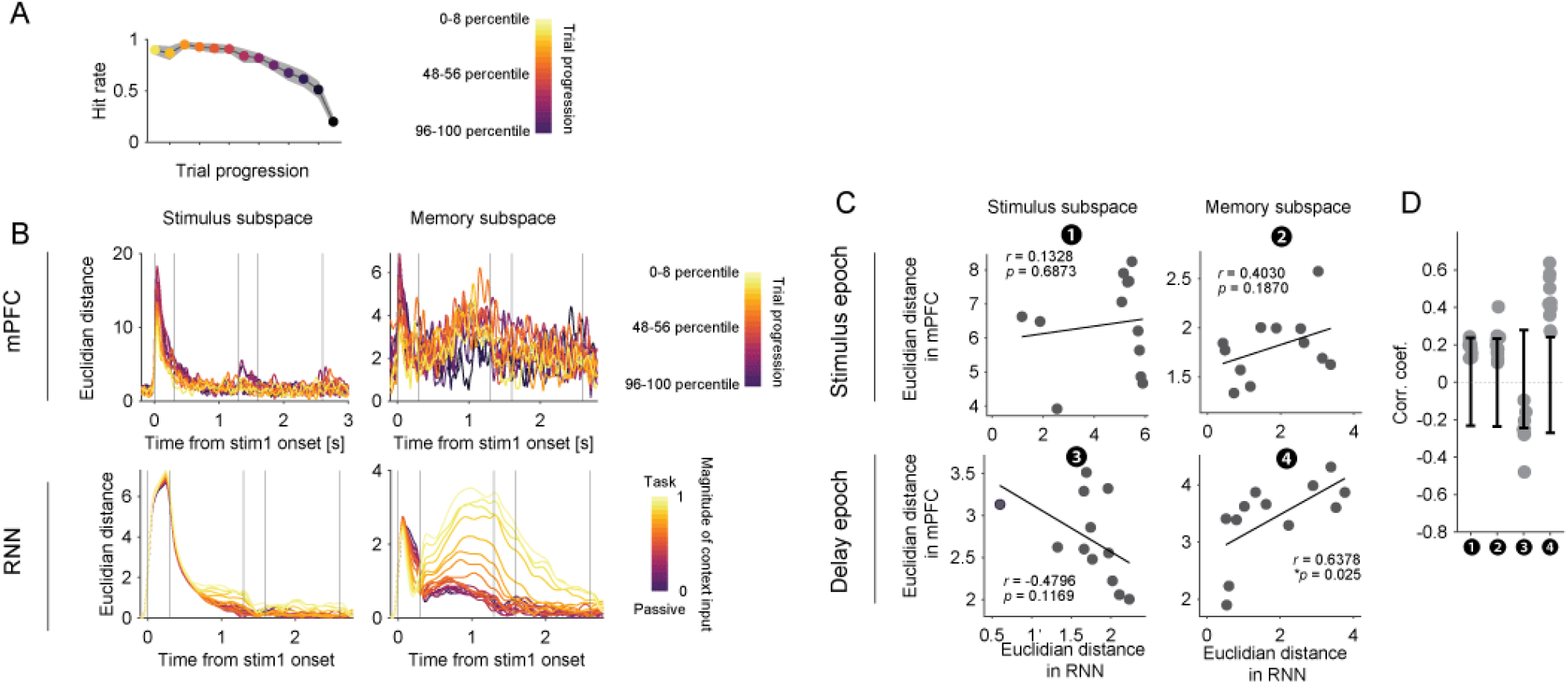
Task motivation correlates with context-input driven fluctuations in memory subspace. **(A)** Behavioral performance (hit rate) of mice as a function of trial progression, reflecting fluctuations in task engagement. **(B)** Euclidean distance between low- and high-tone trajectories in the stimulus and memory subspaces for mPFC across trial progression, and for RNNs with interpolated context input. **(C)** Correlation between fluctuations in Euclidean distance of mPFC and example RNN activity during stimulus and delay epochs in the stimulus and memory subspaces. **(D)** Summary of correlation coefficients between mPFC and RNN distance fluctuations across all trained networks (N = 10). Error bars represent mean ± 2 SD (capturing 95.5% of the variance). Each dot represents an individual network.

### Potential mechanism of dynamic reconfiguration of network dynamics

Similar to the brain, the data-constrained RNN exhibited complex dynamics in a high-dimensional space. This makes it difficult to rigorously characterize the dynamics within the memory subspace. In particular, the system is not reducible to a low-dimensional model that would allow an explicit mapping between the bifurcation structure and the activity of individual units under varying context input. To address this limitation, we constructed a simplified neural field model of interacting neural populations that qualitatively reproduces the bifurcation observed in the data-constrained RNN^36–38^ (**Figure 4B**). We modeled two distinct mechanisms by which context input can modulate the system. First, we considered a bias model in which context input is added linearly to both units (**Figure 6A**). Second, we implemented a gain model in which context input modulates the strength of inhibition between the two units, thereby controlling the effective interaction gain (**Figure 6B**). Previous studies have shown that, in such models, modulation of a shared input or weight strength between units can reconfigure recurrent dynamics, enabling transitions between regimes with a single stable fixed point, multiple fixed points, or dynamics approximating a line attractor^39–41^. By comparing the behavior of these two models with mPFC neuronal data, we sought to gain mechanistic insight into how network dynamical regimes can be flexibly reconfigured by context input.

**Figure 6.**
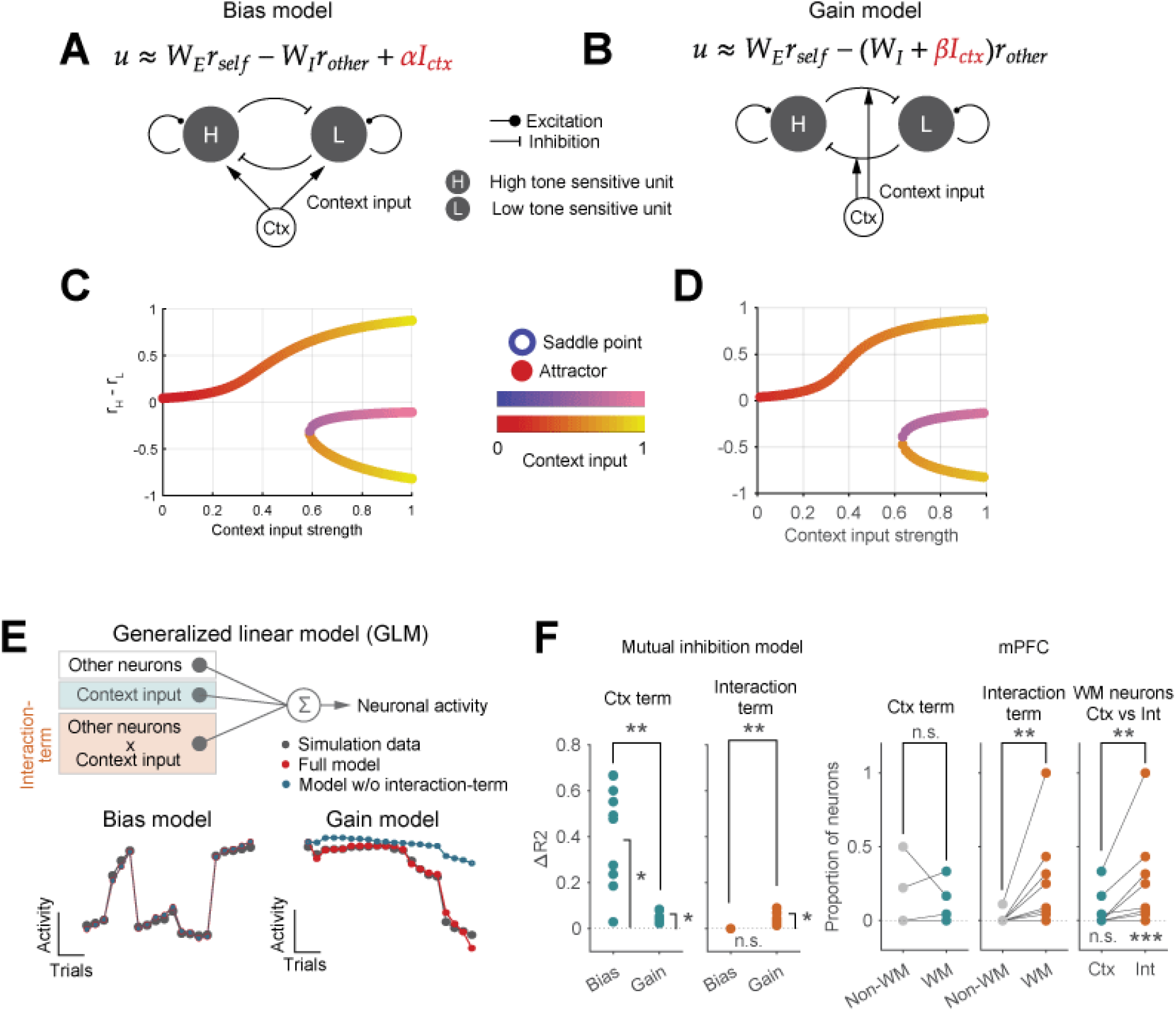
Two distinct types of mutual inhibition model. **(A-B)** Schematics of the mutual inhibition models. Context input provides either an additive drive to both units (Bias model, **A**) or a multiplicative modulation of the strength of mutual inhibition (Gain model, **B**). **(C-D)** Attractors and saddle points during interpolation between contextual input strengths corresponding to passive (ctx = 0) and active (ctx = 1) task conditions for both models. **(E)** (Top) Schematic of the generalized linear model (GLM). The activity of each model unit is regressed on the activity of the other unit, context input, and their interaction term. (Bottom) Example predictions of the GLM with (red) and without (blue) the interaction term for the bias (left) and gain (right) models. Each dot represents a trial (***p < 0.001, two-sample t-test). **(F)** (Left) Change in predictability between the full model and the model without the context or interaction terms. Values near zero indicate a negligible contribution of the interaction term. (Right) Proportion of neurons for which the context and interaction terms significantly improved the model predictability, are shown separately for working memory (WM) and non-WM neurons (Ctx term and interaction term). Each dot represents a session, with lines connecting paired sessions (*p < 0.05, paired t-test).

As in previous studies, systematically varying the strength of the context input led to a bifurcation in both models (**Figure 6C-6D, S6A-B**) similar to the data-constrained RNN (**Figure 4B**). Importantly, stronger context input drove the units into the saturating regime of the nonlinear activation function, where the slope was reduced (**Figure S6C-S6D**). In the bias model, a gradual shift of the operating point toward a shallower slope reduces the effective coupling and stabilizes the activity of the two units, and facilitates their sustained dynamics. In contrast, in the gain model, modulation of context primarily acts through changes in the strength of mutual inhibition, thereby reshaping the stability landscape of the system. As a consequence, the balance between self-excitation and mutual inhibition is shifted, leading to changes in the location and stability of fixed points. Importantly, this modulation does not directly target the single-unit gain function, but instead changes the network-level effective coupling, which can indirectly influence the local operating point through recurrent interactions. In our implementation, this results in both a shift of the fixed-point structure and an apparent change in the local sensitivity of the response (**Figure S6D**), even though the primary mechanism is the modulation of inhibitory coupling strength rather than the nonlinear gain itself.

What differed between the models was how context acted on neural activity. In the bias model, context is an additive bias, while in the gain model, context modulates recurrent connections, leading to a multiplicative effect on neural activity (**Figure 6A-6B**, see Supplementary Notes for details). This difference is reflected in the fact that, in the bias model, neural activity could be well explained as a linear combination of context and the activity of the other neural pool (**Figure 6E-6F**, bias model). While, in the gain model, there was a significant interaction between context and neural activity, reflecting the modulation of connections between the two units (**Figure 6E** and **6F**, gain model). To test these predictions in the brain, we quantified the extent to which mPFC neuronal activity could be explained by the activity of other neurons, context input, and their interaction term. Our previous analysis indicated that context information predominantly influences activity during the working memory (WM) period. Accordingly, we applied the GLM separately to populations of WM-selective and non-WM neurons. Here, the strength of context input was estimated as block-wise hit rate within the task (**Figure 5A**). As described above, hit rate was negatively correlated with task engagement, and was therefore treated as a non-zero, gradual modulation of context input. In the mPFC, the contribution of the interaction term was significantly greater in WM-selective neurons than in non-WM neurons (**Figure 6F**, mPFC). Moreover, in WM neurons, the interaction term contributed significantly more to model performance than the context term alone (**Figure 6F**, Ctx vs Int). These results suggest that context-dependent input selectively modulates memory-related neuronal populations, by altering their interactions, leading to a bifurcation in population dynamics.

## Discussion

In this study, we combined experimental neuronal data from the mouse brain with computational modeling to investigate how context can flexibly modulate the dynamics of neural networks. Using a working memory task, we compared population activity in the mPFC during active task engagement and passive stimulus exposure (**Figure 1**). Switching between active and passive task contexts predominantly affected neural representations in the memory-maintenance subspace (**Figure 1** and **3**). To mechanistically examine how context modulated network dynamics, we fit data-constrained RNNs and systematically interpolated the strength of context inputs (**Figure 2** and **3**). This showed that changing the context shifted the network dynamics from a regime with a single stable attractor to a regime with a bistable attractor, effectively gating memory-related dynamics in a context-dependent manner (**Figure 4**). Importantly, similar results were seen in neural recordings; when the animals’ behavior was more consistent, the separation of memory representations was greater (**Figure 5**). Finally, simulations using the reduced model suggest that context modulates the interactions between neural populations that represent different items in working memory (**Figure 6**). Together, our results suggest that variability in internal state—approximated here by context input—can influence working-memory performance by reshaping the underlying dynamical landscape. In this way, cognitive performance depends not only on task demands but also on internal motivational variables that regulate network dynamics.

Our analyses further revealed how internal states selectively act on specific components of population activity. While stimulus encoding dynamics were preserved across contexts, the dynamics within the memory maintenance subspace were strongly modulated by both task context and behavioral engagement, as indexed by hit rate (**Figure 3A** and **5B**). Moreover, these changes in dynamics are consistent with modulation of effective coupling strength between specific neuronal populations (**Figure 6B** and **6F**). Together, these findings provide a perspective distinct from previously described mechanisms in which additive external inputs shift network dynamical regimes^39,40^, instead highlighting a role for context-dependent modulation of recurrent interactions.

There are at least two biologically plausible mechanisms that could implement such selective modulation of neuronal interactions among specific subpopulations. First, internal state signals may be conveyed by dedicated circuits that preferentially project to subsets of mPFC neurons, particularly inhibitory neurons. These inhibitory neurons may then regulate the effective synaptic interactions among WM neurons^42,43^, thereby modulating their functional coupling. Through this circuit mechanism, internal state signals could dynamically reshape the interaction within the local network, rather than directly altering the synaptic transmission at the level of individual synapses. Consistent with this, fluctuations in pupil diameter—often used as a proxy for arousal state—have been shown to strongly correlate with the activity of specific interneuron populations, including vasoactive intestinal peptide (VIP) and somatostatin (SST) interneurons. Moreover, studies in the dentate gyrus have shown that SST interneurons can regulate winner-take-all dynamics by modulating interactions among excitatory neurons^44^.In addition, interneurons have been reported to modulate neuronal activity through both divisive and subtractive normalization^45^, thereby flexibly controlling the activity of neurons. Second, state– dependent release of neuromodulators could selectively modulate the gain of recurrent interactions^46–48^. For instance, dopamine, acetylcholine, noradrenaline are known to regulate arousal^49–51^, attention^52^, and task engagement^53–55^ and may preferentially influence neurons supporting memory maintenance. This is consistent with prior studies implicating neuromodulators, such as dopamine, in the regulation of working memory^56–59^. How these signals are integrated at the circuit level within mPFC remains an important open question.

The selective modulation of specific subspaces is well aligned with the modular organization of prefrontal cortex. Recent studies suggest that mPFC exhibits a structured modular architecture^34,60^, with distinct subnetworks supporting different cognitive functions. Within this framework, internal state signals could selectively engage particular modules, thereby inducing transitions in the dynamical regime of the network. Linking functional subspaces identified at the population level to underlying circuit architecture may therefore provide a powerful approach for understanding how internal state inputs reshape cortical dynamics.

Our findings extend existing frameworks of attractor dynamics and context-dependent computation. Previous studies have shown that recurrent circuits can reconfigure information selection and the corresponding recurrent dynamics by incorporating contextual input^61–63^. Here, we demonstrate that internal state variables—modeled as continuous context inputs—do not simply switch the network between discrete configurations. Instead, they continuously modulate the geometry of mnemonic attractors in mPFC, including the induction of bifurcations that generate multiple stable states (**Figure 4B** and **4C**). This continuous control of attractor structure provides a natural mechanism by which gradual changes in motivation or engagement can lead to graded changes in working-memory fidelity, without altering stimulus representations themselves (**Figure 5A** and **5B**). Together, these results support a view in which internal states dynamically sculpt attractor landscapes in higher-order cortical areas, providing a principled account of how motivation and engagement modulate cognitive computation.

## Acknowledgements

We thank laboratory members of Sur lab, especially Jung Won Bae and Marco Celotto for fruitful discussion and feedbacks, and Toshitake Asabuki, Yoshihito Saito, Kensuke Yoshida for constructive feedback on this paper. This work was supported by NIH grants R01MH133066 and R01NS130361, MURI Grant W911NF2110328, and The Picower Institute Innovation Fund (M.S.); NIH grants R01MH126022 and R01MH129492 (T.J.B.); Japan Society for the Promotion of Science (JSPS) Overseas Research Fellowships, and The Uehara Memorial Foundation Postdoctoral Fellowship (Y.O.).

## Author Contributions

Conceptualization: Y.O.

Methodology: Y.O.

Software: Y.O.

Validation: Y.O.

Formal analysis: Y.O.

Investigation: Y.O.

Resources: Y.O., M.S.

Data Curation: Y.O.

Writing - Original Draft: Y.O.

Writing - Review & Editing: Y.O., T.J.B., M.S.

Visualization: Y.O.

Supervision: T.J.B., M.S.

Project administration: Y.O.

Funding acquisition: Y.O., T.J.B., M.S.

## Declaration of Interests

The authors declare no competing interests.

## Supplemental figures

**Figure S1.**
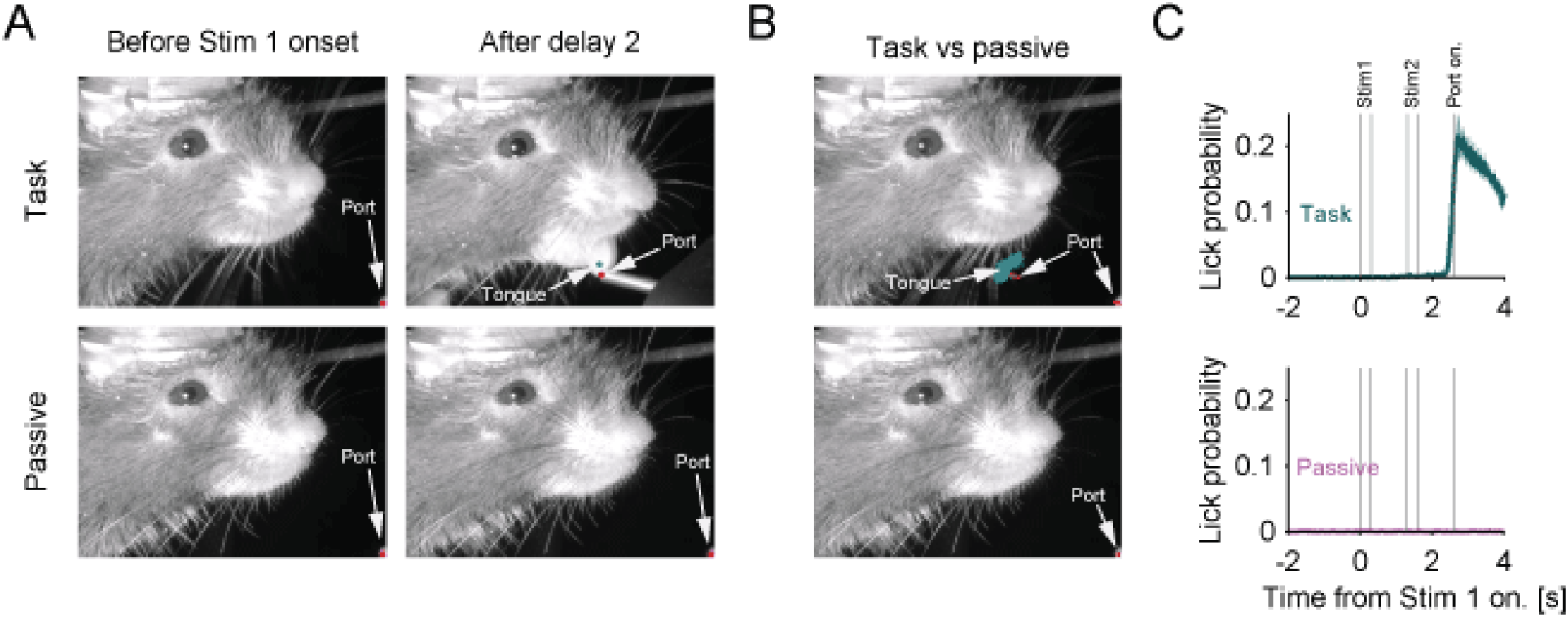
Lick probability during task and passive conditions. **(A)** Example frames of the mouse face during task and passive conditions. Left and right columns correspond to frames acquired before stimulus 1 onset and after delay 2 offset, respectively. **(B)** All licking events and port labels during task and passive conditions. Mice did not exhibit licking behavior during passive conditions. **(C)** Lick probability during task and passive conditions. Lines and shaded areas indicate the mean ± 95% confidence intervals.

**Figure S2.**
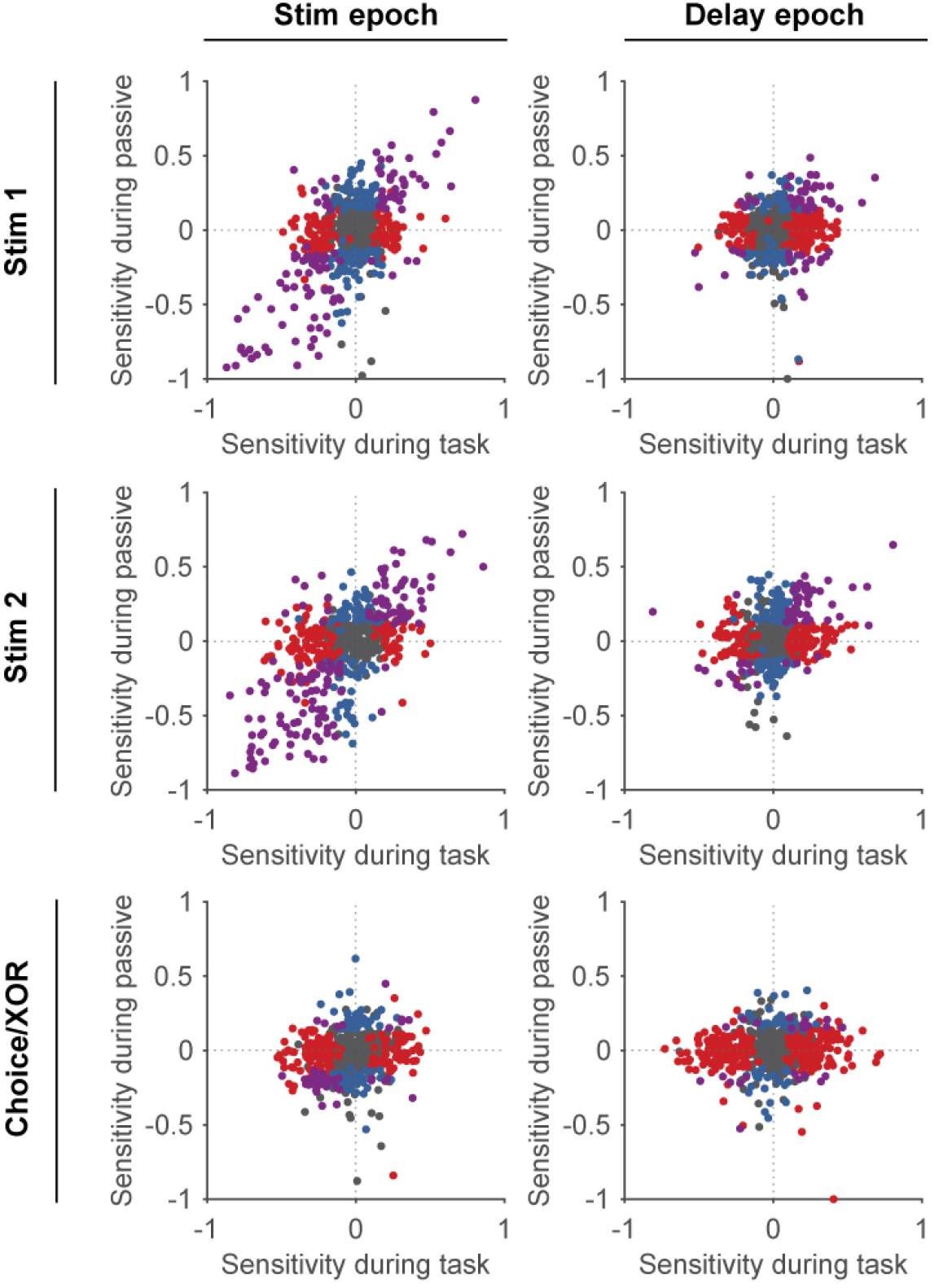
Neuronal selectivity for Stim 1, 2 and choice/XOR. Scatter plots and histograms illustrate stimulus- and choice-related modulation of mPFC neuronal activity during task engagement and passive stimulus presentation. Neurons exhibiting significant selectivity for either the high or low tone during the stimulus epoch in the task condition largely preserved the same tone preference during the passive block (top left; r = 0.5472, p = 9.99 × 10^−4^, permutation test). In contrast, stimulus selectivity observed during the Delay 1 period in the task was only weakly expressed during passive presentation (top right; r = 0.1374, p = 0.0020, permutation test). Similar relationships were found for stimulus 2 (middle row; stim epoch: r=0.5854, *p*=9.99e-4, delay epoch: r=0.2343, *p*=9.99e-4, permutation test). Choice- or XOR-related modulation showed a weak correlation between task and passive conditions during the stimulus epoch (r = 0.1251, p = 0.0080, permutation test), but no significant correspondence during the delay epoch (r = 0.0665, p = 0.1049, permutation test). Colored markers indicate neurons significantly modulated by stimulus or choice in the task condition (red), passive condition (blue), or both (purple).

**Figure S3.**
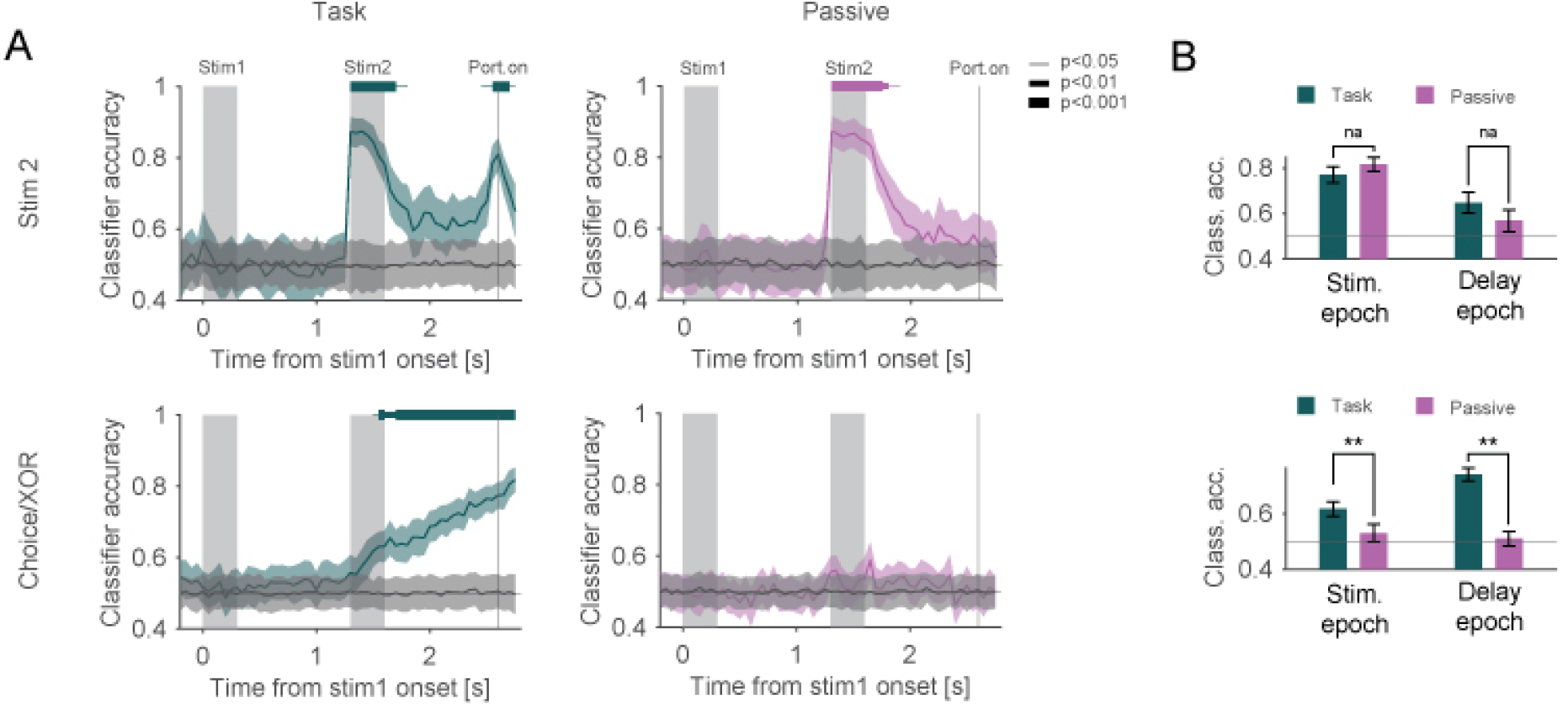
Population analysis for determining whether mPFC and PPC encode stimulus and choice information during active task and passive block. **(A)** Time course of classifier accuracy for stimulus 2 (top row) and choice/XOR (bottom row) in the active task (left) and passive block (right). Horizontal bars above each plot indicate periods of above-chance classification (P < 0.05, 0.01, and 0.001 for thin, medium, and thick lines, respectively; bootstrap test). **(B)** Proportion of neurons encoding stimulus 2 and choice/XOR during stimulus and delay epochs in active and passive tasks. Choice/XOR classification accuracy was significantly reduced in passive block (p < 0.05, bootstrap test).

**Figure S4.**
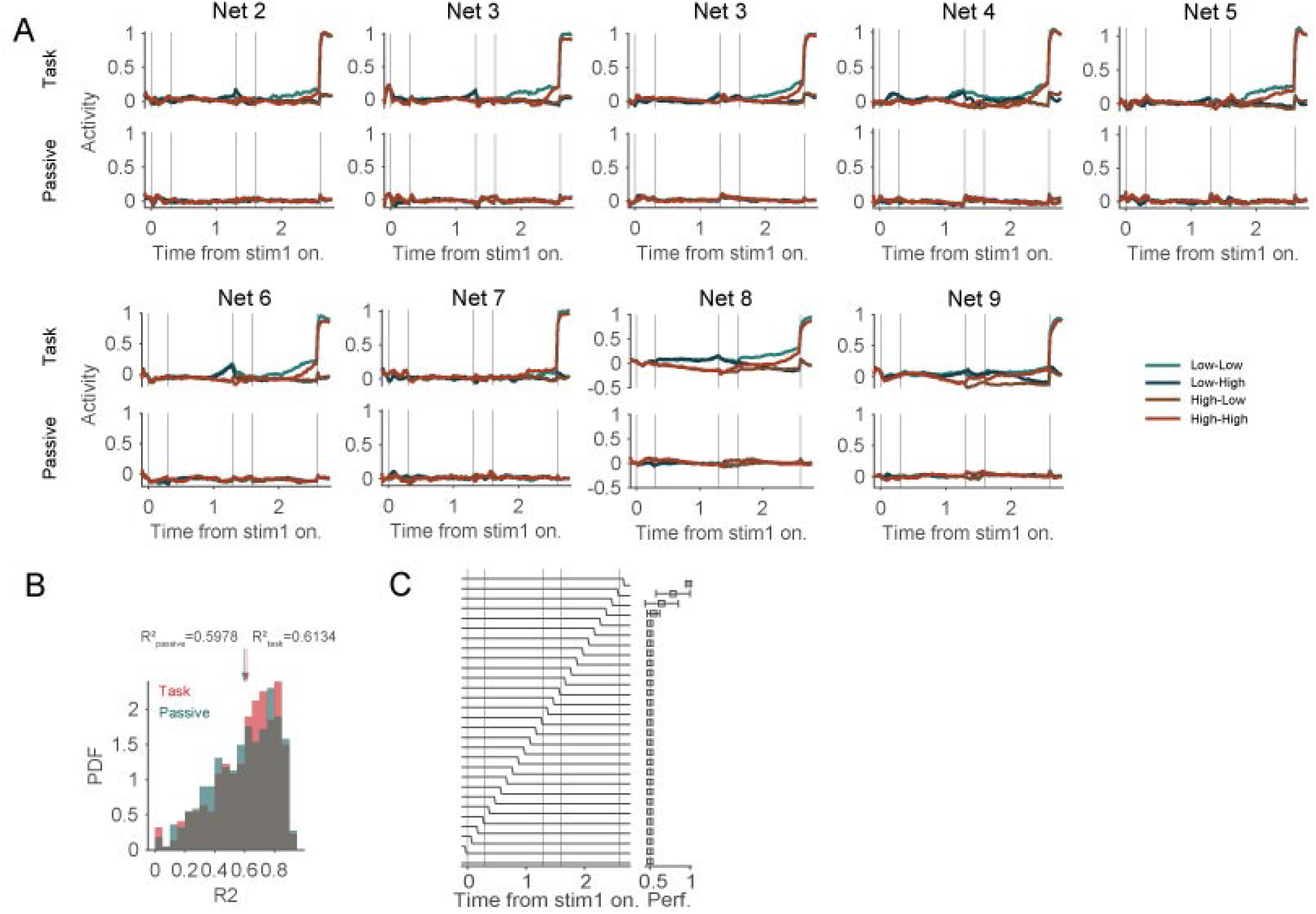
Output activity and R^2^ value in additional data-constrained networks. **(A)** Activity of the output unit across all trial types in the task and passive conditions. **(B)** Probability density functions of the coefficient of determination (*R*^2^) computed from network activity during the task (red) and passive (blue) conditions across all trained networks. Allows indicate the average *R*^2^ values for each condition (*Mean* 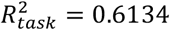 and *Mean* 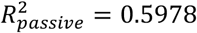). No significant difference was observed between the two distributions (*p* = 1.0, Two-sample Kolmogorov-Smirnov test). **(C)** RNN performance under different temporal profiles of context input. Context input was abruptly reduced from 1 to 0 at varying time points within the trial. The data are shown as mean ± SEM across networks (N=10).

**Figure S5.**
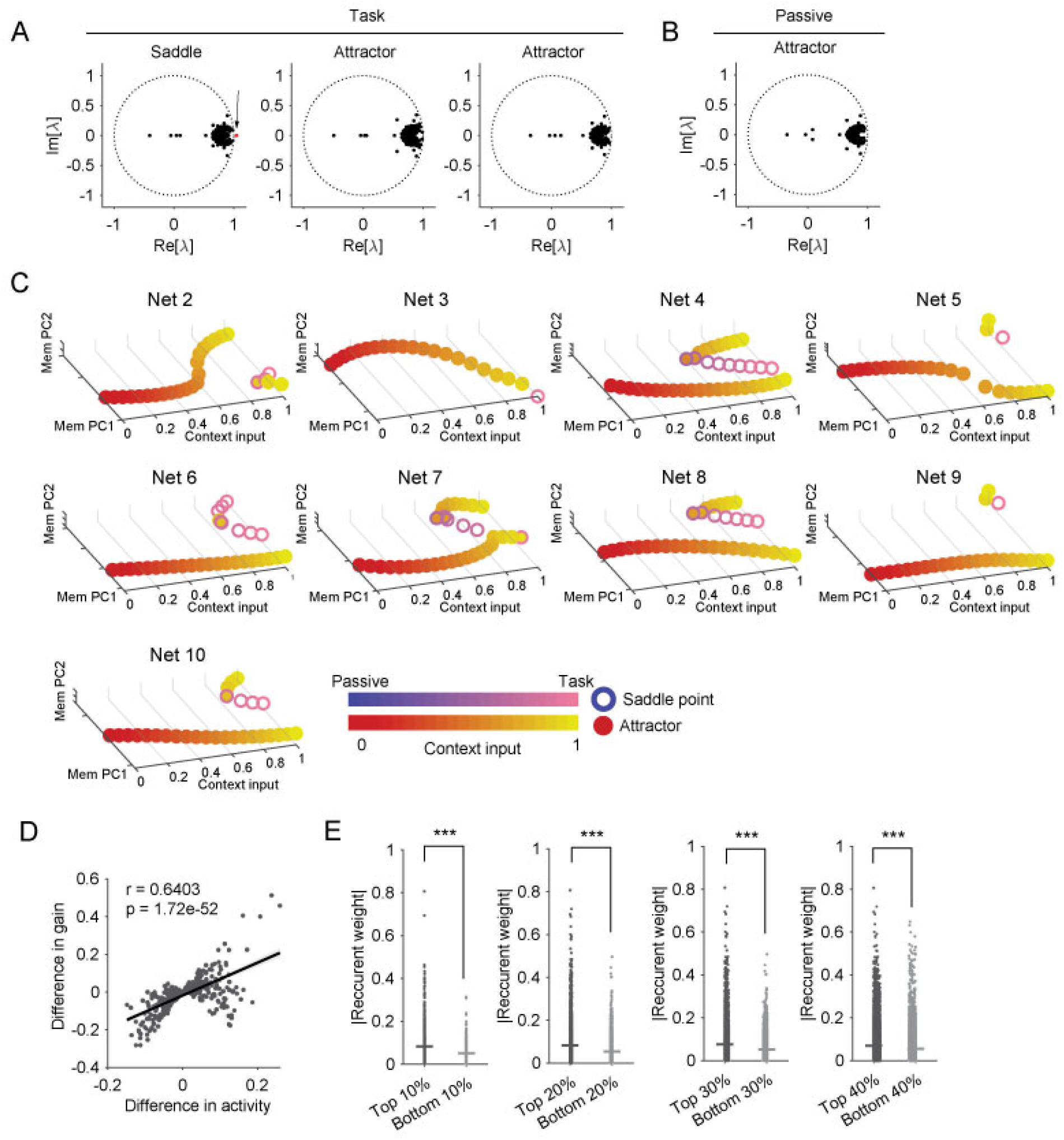
Fixed point structure and bifurcation diagrams in additional networks. **(A)** Eigenvalue spectra in the complex plane for fixed points identified in the task condition from an example network. Black dots indicate eigenvalues of the Jacobian evaluated at each fixed point; the red dot denotes an eigenvalue with magnitude greater than 1, indicating an unstable direction and thus a saddle point. **(B)** Same as (A), but for passive condition. **(C)** Attractors and saddle points during interpolation between contextual input strengths corresponding to the passive (input = 0) and active task (input = 1) conditions, visualized in the memory subspace. Dots and circles denote attractors and saddle points, respectively, with color indicating the magnitude of contextual input. **(D)** Relationship between changes in unit activity and gain between the passive and active task conditions. Each point represents an individual RNN unit. r denotes the Pearson correlation coefficient. The correlation was highly significant (p = 1.72 × 10^−52^, Wilcoxon signed-rank test). **(E)** Absolute recurrent connectivity weights between the top and bottom 10–40% of units exhibiting the largest differences in activity slopes between the passive and active conditions.

**Figure S6.**
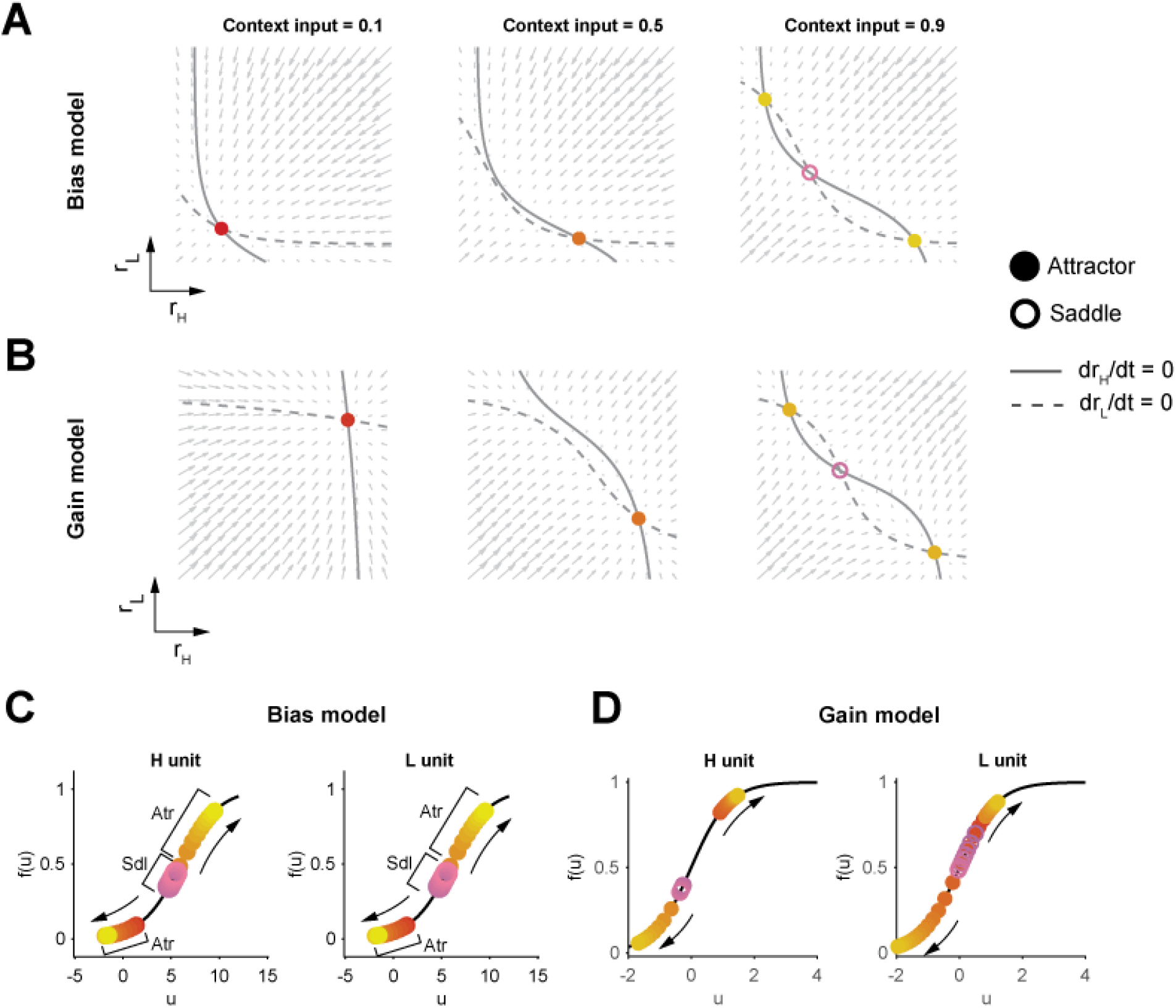
Phase-plane analysis of bias and gain models. **(A-B)** Phase-plane analysis of the bias (**A**) and gain (**B**) models. The two axes represent the activities of the two units (L and H). Context input is applied to both units and drives a reconfiguration of the dynamical landscape. Varying the level of context input shifts the sigmoidal nullclines of the two units (*dr*_*H*_ /*dt* = 0, solid; *dr*_*L*_ /*dt* = 0, dashed line; see Supplementary Notes), thereby altering the number and stability of fixed points. The plots show the nullclines and corresponding fixed points for three levels of context input (0.1, 0.5, and 0.9). (**E-F**) Interaction between context input and the nonlinear (sigmoidal) activation function in both models. Increasing context input shifts the operating regime toward lower effective gain, leading to corresponding changes in nullclines and a reorganization of the fixed-point structure.

## Methods

### Behavioral task and recordings

This experiment has been described in detail previously^34^. Here we provide a brief summary of the behavioral setup and data acquisition. Briefly, male and female mice (>8 weeks old) were housed under a reversed light/dark cycle (lights off 09:00–21:00) with controlled temperature and humidity, and all experiments were performed during the dark phase. All procedures were approved by the Massachusetts Institute of Technology’s Animal Care and Use Committee and adhered to NIH guidelines.

Mice were head-fixed in a custom behavioral rig and trained on a DMS-dr task with auditory stimuli. On each trial, white LED is presented to indicate the beginning of each trial. After a 1-1.5s, two pure tones (3 or 14 kHz, 0.3 s duration) were presented sequentially with a 1s delay between them and followed by second 1s delay. Mice learned to lick a spout for water reward when the tones matched, and to withhold licking when the tones differed. Incorrect responses were punished with a siren-like auditory stimuli, air-puff, and extra 7.0s inter-trial-interval.

For in vivo electrophysiology, trained mice were anesthetized with isoflurane for craniotomy over mPFC a day before the recording. NeuroPixels 1.0 probes^64^ (IMEC) were inserted at controlled angles and depths, and recordings were acquired at 30 kHz. Spike sorting was done using Kilosirt2.5^65^, and spikes were manually curated using Phy GUI (https://github.com/kwikteam/phy) to remove artifacts. Units with an inter-spike interval (ISI) violation ≤ 0.5, firing rate ≤ 0.1 Hz, or ill-shaped waveform were filtered out. Spike times were verified with cross correlograms to combine units or eliminate duplicates. For each unit, part of the recordings with obvious drift (unit spikes abruptly disappearing) were excluded. A total of 586 neurons were recorded from 18 sessions that included 6 different mice.

### Videographic analysis

We used a machine-learning-based pose estimation algorithm (DeepLabCut^66^ v3.0.0) to automatically track the licking port and tongue movements in behavioral videos acquired during recordings (**Figure S1**). A subset of video frames (40–50 frames) was manually annotated to label the positions of the licking port and tongue for network training. After training, the x- and y-coordinates of the port and tongue were automatically extracted from each video frame.

### Lick probability

To quantify the task motivation in mice, we calculated the lick probability (hit rate) in successive blocks of trials (**Figure 5A**). For each session, trials were divided into percentile-based blocks with an 8% step size across the progression of training. Within each block, the hit rate was computed as the proportion of correct licked response among all matched trials: *Hit rate* = *N*_*hit*_ /(*N*_h*it*_ + *N*_m*i*ss_), where *N*_h*it*_ and *N*_m*i*ss_ denote the number of hit and miss trials, respectively.

Hit rates were first calculated independently for each session and then averaged across sessions. To assess whether hit rates systematically changed across training progression, we computed the Pearson correlation coefficient between block order and the mean hit rate across blocks.

### Analysis of neural activity

For all mPFC neurons, changes in firing rate associated with behavior were assessed using peri-stimulus time histograms (PSTHs). Unless otherwise stated, PSTHs were computed using 10 ms bins for individual neurons in each recording session and smoothed with a Gaussian filter with a standard deviation of 20 ms to obtain the temporal profile. For visualization and analysis, firing rates were z-scored across sessions.

### Selectivity

All comparative indices (stimulus 1, 2, and choice selectivity index) were computed using a receiver operating characteristic (ROC) analysis, which calculates the ability of an ideal observer to classify whether a given spike density was recorded under one of two trial types (**Figure 1D** and **S2**). We indexed the difference between two firing rate distributions by scaling the ROC area between -1 and 1, where 0 reflects no difference between the distributions and the sign denotes whether a neuron fires more under one of the trial types than the other. Statistical significance (p<0.05) was determined with a permutation test of 500 repetitions. In this analysis, we used firing rates in only correct trials during the task or all trials in passive task.

We assessed stimulus 1 and 2 selectivity by comparing high and low tone trials at the stimulus or delay period (stimulus period: 0-0.5s for stimulus 1 and 1.3-1.8s for stimulus 2, delay period: 0.8-1.3s for stimulus 1 and 2.1-2.6s for choice/XOR relative to stimulus 1 onset). Choice selectivity was computed by comparing lick and non-lick trials at the delay period during the active task or comparing matched and non-matched trials during passive block.

### Classification analysis

For classification analysis (**Figures 1E** and **S3A**), we used neurons that were recorded in the sessions with >65% correct rate. For classifiers, we used a logistic regression classifier as implemented by the MATLAB *fitclinear* function to quantify the amount of information about the stimulus (low or high tone) and choice/XOR (lick or non-lick trial and matched or non-matched trial) in the population of neurons. This analysis included all neurons that were recorded during at least 30 trials for each stimulus and choice/XOR trial types, and we constructed “pseudo-trials” by randomly extracting trials from desired conditions for each neuron^67^. For the training and testing dataset, the number of trials in each condition was matched to prevent bias in training classifiers. We also randomly chose the trials of lick trials from low-low and high-high trials and non-lick trials from low-high and high-low trials with equal probability (15 trials for each combination). Classifiers with L2 regularization (λ = 1/20) was trained to classify the stimulus or choice using pseudo-population data and tested on held-out data. The choice of the λ value did not affect our conclusion. We used tenfold cross-validation by leaving a 10% subset of trials for classification to avoid overfitting. This procedure was repeated 100 times. Classification results are reported either as a function of time. Time-resolved classification was done on z-scored firing rates measured in a 50 ms moving window. Statistical significance was tested by comparing the resampling distribution of classification accuracy averaged within tenfold cross-validation against classification accuracy of shuffled label classifier. If these distributions are not overlapped in a 2SD range (95.5% data in this range), they are defined as significantly dissociated.

### Generalized linear model

To examine whether task motivation modulates functional interactions between neurons, we applied generalized linear modeling (GLM) to simultaneously recorded neuronal populations (**Figure 6F**). Sessions with behavioral performance below 65% correct were excluded from the analysis.

Neurons were classified as working memory (WM) or non-WM neurons based on a stimulus selectivity index computed during the late delay 1 epoch (0.8–1.3 s after stimulus 1 onset), as described above. Neurons with a selectivity index exceeding the threshold corresponding to P < 0.1 were classified as WM neurons, whereas the remaining neurons were classified as non-WM neurons.

To characterize gradual changes of task motivation across the session, trials were divided into percentile-based bins using 8% increments, and the hit rate was calculated for each bin, as described above. Neuronal activity from all trials within each bin was pooled, and the corresponding normalized hit rate was assigned to each trial. For each neuron, activity was predicted from the activity of all simultaneously recorded neurons, the normalized hit rate, and interaction terms between neuronal activity and normalized hit rate according to the following model:

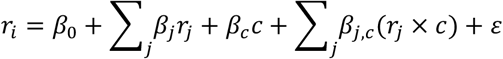

where *r*_*i*_ denotes the activity of the target neuron, *r*_*j*_ represents the activity of simultaneously recorded neurons, *c* corresponds to the normalized hit rate (referred to as internal state variable), *r*_*j*_ × *c* denotes the interaction terms between neuronal activity and hit rate, *ε* represents residual error. *β*_0_ , *β*_*j*_ , *β*_*c*_ , and *β*_*j*,*c*_ denote the intercept, neuronal coefficients, hit-rate coefficient, and interaction coefficients, respectively.

Predictors and target activity were z-scored before model fitting. Models were fit using L1-regularized linear regression (Lasso, *λ* = 0.001). Prediction performance was evaluated using 15-fold cross-validation. To quantify the contribution of the interaction terms, model performance (*R*^2^) was compared between the full model and a reduced model in which the interaction coefficients were set to zero during prediction. Statistical significance for each neuron was assessed using one-tailed t-tests across cross-validation folds followed by Holm–Bonferroni correction for multiple comparisons. For each session, we calculated the proportion of neurons showing significant interaction contributions separately for WM and non-WM populations. Group-level comparisons between neuronal populations were performed using paired t-tests.

### Data constrained RNNs

We constructed a rate-based RNN in which each unit was trained to reproduce the peri-stimulus time histogram (PSTH) of a single experimentally recorded neuron from the mouse medial prefrontal cortex (mPFC), pooled across recording sessions^68,69^ (**Figure 2**). All networks are a time-discretized RNN with positive activity. Before time discretization, the network state *h*(*t*) evolves according to a dynamical equation:

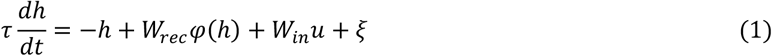

Here, *τ* denotes the units’ time constant, *W*_*rec*_ and *W*_*in*_ are the recurrent and input weight matrices, respectively, *u* represents the external input vector, and *ξ* indicates additive Gaussian noise.

The nonlinear activation function φ(·) was defined as a sigmoid with a gain parameter *β* = 3.0.

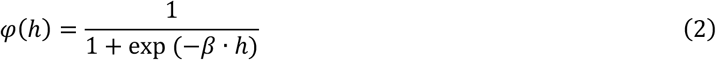

Network output unit *z* was read out from the network according to:

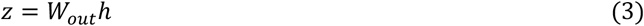

where *W*_*out*_ denotes the output weight matrix of size *N*_*out*_ × *N*_*rec*_ . All weight matrices (*W*_*win*_ , *W*_*rec*_ , and *W*_*out*_) were optimized during training.

After using the first-order Euler approximation with a time-discretization step Δt = 20ms, we have

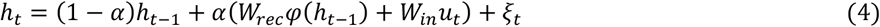

Here, *α* ≡ Δt/*τ*, and we use a discretization step Δt = 20ms. We imposed no constraint on the sign or the structure of the weight matrices *W*_*in*_ , *W*_*rec*_ , *W*_*out*_ . The network and the training are implemented in PyTorch.

The network received four types of noisy input:

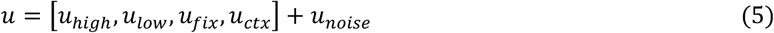

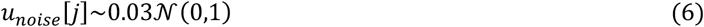

The fixation input *u*_*fix*_ was 1 when the network was required to remain still and 0 when the network was required to respond. The stimulus inputs (*u*_*high*_ and *u*_*low*_) represented high and low tones in the mouse’s experiment, respectively, and were set to 1 during the stimulus epoch and 0 otherwise. The context input (*u*_*ctx*_) was 1 when the network was required to perform DMS-dr task and 0 when the network was required to ignore. The time constant of neurons was *τ* = 100*ms*.

### Task and performance

We implemented an abstraction of the DMS-dr task that mice performed in this study. The task consisted of six periods; initial delay (100 ms), stimulus 1 (300 ms), delay 1 (1000 ms), stimulus 2 (300 ms), delay 2 (1000 ms), and response period (200 ms). Inputs and output for an example trial are shown in Figure 3A. Fixation input remained 1 throughout the trial until the response period, where it changed to 0. Fixation input reflect the suppression of licking behavior in the mouse’s experiment. The average of response output during the response period determined whether the trial was reported as a match (output > 0.5) or non-match (output ≤ 0.5), reflecting licking or non-licking behavior in the mouse’s experiment. The task performance was computed by comparing the response output with successfully reported with match and non-match trials. During passive task condition (*u*_*ctx*_ = 0), network had to produce zero response output.

### Network training procedure

We used back-propagation through time^70^ to train networks to minimize loss functions ℒ. Trials were generated stochastically, following a predefined temporal structure and mapping stimulus inputs to target outputs 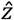 . The total loss function was:

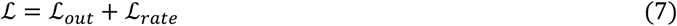

where

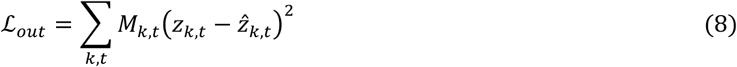

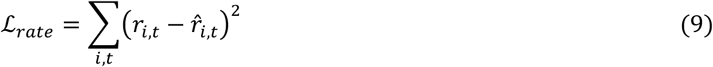

Here, *z*_*k*,*t*_ and 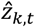 are, respectively, the actual and the target readout values and the indices *kk* and *t* , respectively, are the index of the output and timesteps. *r*_*i*,*t*_ and 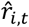 are, respectively, the actual and the trial-averaged mPFC neuronal activity of the *i*-th neuron. We implemented a mask, *M*_*k*,*t*_ , for modulating the loss with respect to certain time intervals.

- For the response output unit (*k* = 1), *M*_1,*t*_ = 0 before stimulus 1 onset, *M*_1,*t*_ = 5 during the response period, and *M*_1,*t*_ = 1 for the rest of the trial.
- For the fixation output unit (*k* = 2), *M*_2,*t*_ = 0 before stimulus 1 onset and *M*_2,*t*_ = 2 for all other timepoints. We trained networks with *N* = 443 units to reproduce neural activity recorded from mPFC during active and passive tasks. The networks were trained using Adam optimizer^71^ in PyTorch with the decay rate for the first and second moment estimates of 0.9 and 0.999 and learning rates of 10^−2^, respectively. During training, we used mini-batches of 32 trials. In a given minibatch, all trial types were randomly generated. Training was terminated when ℒ_*out*_ fell below 0.02, and the task performance exceeded over 0.99.

### Fixed point analysis

To characterize the underlying dynamical structure of the trained RNN (**Figures 4A-C**, and **S5A-C**), we searched for fixed points of the network dynamics following the approach of Sussillo and Barak^62,72^.

Fixed points *h*^*^ satisfy

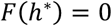

where

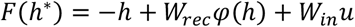

and φ(·) is the sigmoid activation function. To identify such points numerically, we define the scalar “speed” function

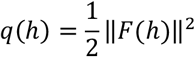

which is zero exactly at fixed points. We minimized *q*(*h*) using gradient-based optimization, where the gradient is given by

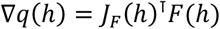

where *J*_*F*_ (*h*) is the Jacobian of *F* with respect to *h*.

Optimization was repeated from multiple random initial conditions drawn uniformly from the state space. We used MATLAB *fminunc* function with analytically provided gradients. Candidate solutions whose final speed satisfied *q*(*h*) < 10^−3^ were retained as valid fixed points.

Because repeated optimizations from different initial conditions can converge to the same fixed point, we removed duplicate solutions by clustering. Pairwise Euclidean distances between all candidate fixed points were computed, followed by hierarchical clustering using single-linkage. Fixed points within a distance threshold of 10^−3^ were clustered together, and a single representative from each cluster was retained for subsequent analyses.

To classify the stability of each fixed point, we analyzed the local linearized dynamics around each fixed point. Rather than using the continuous-time Jacobian eigenvalues directly, we assessed stability using a discrete-time approximation of the dynamics, consistent with the numerical integration used in simulations. Specifically, we computed the Jacobian *J*_*F*_ (*h*^*^) at each fixed point and constructed the discrete-time Jacobian

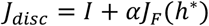

where *I* is the identity matrix and *α* = 0.2 is the effective integration step size. We then computed the eigenvalues {*λ*} of *J*_*disc*_.

Eigenvalues were classified as stable if |*λ*| < 0 and unstable if |*λ*| > 0. Based on this criterion, fixed points were classified as follows:

- Attractor: all eigenvalues are stable
- Saddle: both stable and unstable eigenvalues are present
- Repeller: all eigenvalues are unstable

### Correlation between RNN and mPFC activity

To compare the neuronal dynamics between mPFC activity and RNNs, we computed the similarity of population activity in the stimulus and memory maintenance subspaces (**Figure 5C-5D**). For the neural data, population activity was extracted from two task epochs: the stimulus epoch (0–0.5 s after stimulus onset) and the delay epoch (0.8–1.3 s). For each neuron, mean activity was computed separately for the four trial types (Low-Low, Low-High, High-Low, and High-High) using only correct task trials. Population response matrices were then constructed for the stimulus and delay epochs independently. PCA was applied separately to the stimulus-epoch and delay-epoch population activity to define stimulus and memory maintenance subspaces, respectively. Trial-averaged population activity for Low and High stimulus 1 trials was projected onto each subspace, and the Euclidean distance between the Low and High trajectories was computed in each space using the first *n* principal components:

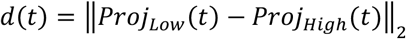

Where *Proj*_*Low*_(*t*) and *Proj*_*High*_(*t*) denote the projected population activity trajectories at time *t*. This analysis yielded time-resolved stimulus representation within the stimulus and memory subspaces.

To examine how representational geometry evolved during learning, task trials within each session were divided into percentile-based training bins using 8% increments across trial progression as described above. Population activity was averaged within each bin before projection into the corresponding subspace. For the RNN, the PCA was applied to unit activity during either the stimulus period or delay period to define stimulus- and memory maintenance subspaces, analogous to the neural analyses. Unit activity was then projected into these subspaces, and Euclidean distances between High- and Low-stimulus trajectories were calculated over time.

To compare population dynamics between mPFC and RNNs, the distance trajectories *d*(*t*) from the RNNs were temporally aligned to the experimental training bins and averaged within corresponding epochs. Pearson correlation coefficients were then computed between the neural distances and the RNN distances for each subspace and epoch. Statistical significance was assessed using a permutation-based null distribution generated by shuffled correlations.

### Mutual inhibition model

Below is a summary of the mutual inhibition models. For detailed descriptions of the models, see the Supplementary Notes.

We constructed a two-unit model in which the two units mutually inhibit each other while each unit possesses self-excitatory recurrent connections (**Figure 6A-6B**). We examined two distinct forms of this mutual inhibition model. In the first model, context input was additively applied to both units. In the second model, context input modulated the mutual inhibition between the two units. Throughout this study, we refer to the former as the “bias model” and the latter as the “gain model”.

### Generalized linear model

To examine how context input modulates unit activity in the mutual inhibition model (**Figure 6E**), we systematically varied the context parameter *c* from 0 to 1 by 0.01 steps and simulated network dynamics for both High-tone and Low-tone conditions. For each context value, the mean activity of the two units during the delay period was calculated.

To determine whether contextual modulation altered the interaction structure between the two units, we fitted generalized linear models (GLMs) to predict the activity of one unit from the activity of the other unit, the context input, and their interaction term. Specifically, the activity of unit 1 (*r*_1_) was modeled as:

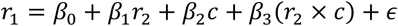

where *r*_2_ denotes the activity of unit 2, *c* represents the context input, *r*_2_ × *c* corresponds to the interaction term, and *ϵ* represents residual error. All predictors were z-scored before model fitting.

Models were fit using L1-regularized linear regression (Lasso) with a linear link function. Prediction performance was evaluated using 10-fold cross-validation. To quantify the contribution of the interaction term, model performance (*R*^2^) was compared between the full model and reduced models in which either the interaction coefficient or the context coefficient was set to zero during prediction. The decrease in cross-validated *R*^2^ was used as a measure of the contribution of each term.

## Data and Code availability

All data and code supporting the findings from this study are available upon reasonable request from the corresponding author (M.S.).

## Notes

### Competing Interest Statement

The authors have declared no competing interest.

