## Supplemental Notes for "Internal state dynamically gates task-specific attractor dynamics in prefrontal cortex"

### Supplementary Notes
#### Mutual inhibition model
#### Network Architecture and Dynamics
To model the network dynamics underlying context-dependent behavior, we constructed a mutual inhibition model consisting of two mutually inhibiting neuronal populations (Unit 1 and 2). The network dynamics are governed by the following standard ordinary differential equations:

$$\tau \frac{dx_H}{dt} = -x_H + F(u_H)$$
$$\tau \frac{dx_L}{dt} = -x_L + F(u_L)$$

where $x_H$ and $x_L$ represent the activation of the two populations, and $\tau$ is the intrinsic time constant of the network. The total synaptic input $u_i$ to each population $i$ is transformed into an activation through a non-linear sigmoidal activation function $F(u)$:

$$F(u) = \frac{r_{max}}{1 + \exp\left(-\frac{u - \theta}{k}\right)}$$

where $r_{max}$ is the maximum activation, $\theta$ determines the threshold (the inflection point of the sigmoid), and $k$ controls the steepness of the activation function.

#### Synaptic currents
The total input variables $u_H$ and $u_L$ integrate recurrent self-excitation, mutual inhibition, baseline inputs, and external stimulus input:

$$u_H = w_{self} x_H - w_{L\to H} x_L + I_0 + I_{bias} + I_{stim(H)}(t)$$
$$u_L = w_{self} x_L - w_{H\to L} x_H + I_0 + I_{bias} + I_{stim(L)}(t)$$

Here, $w_{self}$ is the synaptic weight for recurrent self-excitation, and $I_0$ is a constant background input. To account for intrinsic network asymmetries observed in data-constrained RNN, we introduced a small static bias parameter $I_{bias}$, which slightly favors the activation of H unit over L unit. $I_{stim(i)}(t)$ represents the transient external stimulus received during the task.

#### Alternative mechanisms for contextual gating: bias vs. gain modulation
To elucidate the specific biophysical mechanisms by which the context input reconfigures the attractor landscape, we constructed and compared two distinct mutual inhibition models: a bias model and a gain model. While both models consist of two mutually inhibiting neuronal populations ($x_H$ and $x_L$) and successfully reproduce the context-dependent imperfect bifurcation, they achieve this through fundamentally different mathematical principles, specifically distinguished by the presence or absence of an interaction term between the context and the network state.

##### Bias model
In the bias model, the synaptic weights for self-excitation ($w_{self}$) and mutual inhibition ($w_{L\to H}$ and $w_{H\to L}$) remain constant. The context input ($c$) acts as an external excitatory drive ($I_{ext}$) that is additively applied to both neuronal populations:

$$u_H = w_{self,H} x_H - w_{L\to H} x_L + I_0 + I_{bias,H} + I_{stim(H)}(t) + g_{bias} c$$
$$u_L = w_{self,L} x_L - w_{H\to L} x_H + I_0 + I_{bias,L} + I_{stim(L)}(t) + g_{bias} c$$

where $g_{bias}$ represents the strength of the contextual drive. Mathematically, the context input $c$ is lineally added to the input and is independent of the current state of the network ($x_H$ and $x_L$). The core mechanism of this model relies entirely on the nonlinearity of the sigmoidal activation function $F(u)$. When the context input is low ($c \approx 0$), the additive drive is small, and the network operates in the lower, flat regime of the sigmoid where the effective network gain is insufficient to sustain memory. As the context input increases, the additive drive shifts the operating point of the neurons into the steep, high-gain regime of the sigmoid. In this regime, the constant recurrent weights are effectively amplified, triggering an imperfect bifurcation into a bistable, winner-take-all state. Thus, the bias model gates memory by dynamically shifting the operating point along the nonlinear activation curve.

**Gain model**

In contrast, the gain model assumes that the context input directly alters the effective connectivity of the network. Specifically, the context input $c$ multiplicatively modulates the strength of the mutual inhibition ($w_{L\to H}$ and $w_{H\to L}$) between the populations:

$$w_I(c) = w_{I0} + g_{gain}c$$

where $w_{I0}$ is the original mutual inhibition strength (either $w_{L\to H}$ or $w_{H\to L}$), and $g_{gain}$ is the gain factor that scales the effect of the context. When we substitute this modulated weight into the total input equation for H unit, the mathematical distinction becomes evident:

$$u_H = w_{self,H}x_H - (w_{L\to H} + g_{gain}c)x_L + I_0 + I_{bias,H} + I_{stim(H)}(t)$$

$$u_H = w_{self,H}x_H - w_{L\to H}x_L - \boldsymbol{g_{gain}cx_L} + I_0 + I_{bias,H} + I_{stim(H)}(t)$$

By expanding the equation, a mathematical interaction term ($-g_{gain}cx_L$) emerges. This term dictates that the inhibitory effect of the context input is dynamically scaled by the instantaneous activation of the competing population $x_L$. Therefore, the context input directly stiffens the competitive landscape (effective connectivity) rather than simply shifting the baseline operating point. At low context, mutual inhibition is too weak $w_{I0}$ to separate the states. As the context increases, this state-dependent interaction term forces the network to split into two mutually exclusive memory states through an imperfect bifurcation.

**Parameters**

| Parameters | Bais model | Gain model |
|---|---|---|
| $w_{self,H}$, $w_{self,L}$ | 3.1, 2.9 | 1.5 |
| $w_{L\to H}$, $w_{H\to L}$ | 10, 10 | 0.2, 0.2 |
| $I_0$ | 2.2, -2.2 | 0.1, 0.1 |
| $I_{bias,H}$, $I_{bias,L}$ | 0.2, -0.2 | 0.08, -0.08 |
| $g_{bias}$ | 5.4 | 2.0 |
| $k$ | 2 | 0.6 |
| $\theta$ | 6.0 | 0.0 |
| $r_{max}$ | 1.0 | 1.0 |

**Summary of distinct mechanism**

In summary, the key distinction between the two models lies in whether the context input appears as an additive input term or as an interaction term:

- Bias model contains a context input term ($\boldsymbol{g_{bias}c}$), but no interaction term
- Gain model contains an interaction term ($\boldsymbol{g_{gain}cx_L}$), but no context input term

This distinction reflects two fundamentally different biological mechanisms. The bias model corresponds to non-specific external drive shifting the baseline activity, whereas the gain model corresponds to context-dependent modulation of effective connectivity, potentially mediated by local inhibitory circuits.
